# Perceptual expectations and false percepts generate stimulus-specific activity in distinct layers of the early visual cortex

**DOI:** 10.1101/2022.04.13.488155

**Authors:** Joost Haarsma, Narin Deveci, Nadège Corbin, Martina F. Callaghan, Peter Kok

## Abstract

Perception has been proposed to result from the integration of feedforward sensory signals with internally generated feedback signals. The latter are believed to play an important role in driving false percepts, i.e., seeing things that are not actually there. Feedforward and feedback influences on perception can be studied using layer-specific fMRI, which we used here to interrogate neural activity underlying high confidence false percepts while healthy participants (N=25) performed a perceptual orientation discrimination task. Orientation-specific BOLD activity in the deep and superficial layers of V2 reflected perceptual expectations induced by predictive auditory cues. However, these expectations did not influence participants’ perception. Instead, high confidence false percepts were reflected by orientation-specific activity in the middle input layers of V2, suggesting a feedforward signal contributing to false percepts. The prevalence of high confidence false percepts was related to everyday hallucination severity in a separate online sample (N=100), suggesting a possible link with abnormal perceptual experiences. These results reveal a feedforward mechanism underlying false percepts, reflected by spontaneous stimulus-like activity in the input layers of the visual cortex, independent of top-down perceptual expectations.

## Introduction

Our perception is not always veridical and can become distorted or biased in a myriad of ways, leading to false perceptual inferences. However, the neural mechanisms underlying such false inferences remain hotly debated, with some theories highlighting the contribution of top-down influences such as perceptual expectations, whilst others have pointed out a possible role for feedforward mechanisms. Elucidating the various ways through which perception can go awry is crucial, as it sheds new light on false perceptual inferences as seen in psychiatric disorders like psychosis, and neurological disorders like Parkinson’s disease^1^.

In recent decades there has been an increasing interest in the way our perceptual expectations bias how we interpret the world, which have been proposed to cause false inferences if they influence perception too strongly^2–5^. One particularly popular approach towards understanding such false inferences is predictive coding^5^. According to predictive coding theories, the brain models the world by continuously deploying perceptual expectations regarding the most likely stimuli^6,7^. These perceptual expectations are fed back to the early sensory cortices where they are integrated with sensory inputs to form a percept. The more strongly expectations are weighted, the more they will influence perception, driving it away from sensory input. Therefore, a false percept can arise when perceptual expectations become excessively overweighted^2–5,8^. Indeed, there are many studies demonstrating that expectations play an integral role in shaping sensory experience^9^. Furthermore, there is increasing evidence that individuals experiencing hallucinations demonstrate a stronger influence of perceptual expectations on perception^10–18^.

However, it remains unclear whether all false percepts arise from top-down influences, or whether they can arise from feedforward mechanisms as well. Answering this question is critical as it may shed new light on the different potential pathways towards hallucinogenesis. In line with this, hallucinations in some neurological disorders have indeed been known to arise from fluctuations in feedforward sensory signals^19,20^. For example, in Charles Bonnet syndrome, the visual cortex is partially deafferented, which causes it to become hyperexcitable^4,20–22^. Recent studies of this syndrome suggest that spontaneous local activity emerges in early visual cortex and is fed up the cortical hierarchy, eventually generating a hallucinatory percept^19,22^. This demonstrates that false percepts can at least in principle emerge from feedforward processes. Further, some studies in psychosis have reported increased reliance on sensory evidence, particularly with relation to delusions^15,16,23–27^. Additionally, there is evidence that spontaneous activity can contribute to the experience of false percepts in healthy individuals as well, although it is at present unclear whether these reflect feedforward or feedback signals^28–32^. Here, we used layer-specific fMRI to investigate which cortical layers were involved in representing false percepts.

Sensory feedforward and feedback signals are separated on a laminar level, as they preferentially terminate in different cortical layers. That is, feedback neurons terminate in the agranular deep (layers V and VI) and superficial layers (layers I, II and III), whereas feedforward connections preferentially terminate in the middle granular layers (layer IV)^33,34^. It has recently become possible to non-invasively measure the activity in different cortical layers in humans using layer-specific fMRI, allowing testing of theories about the contributions of feedforward and feedback signals in cognition and perception^8,35–41^.

In the present study, the contribution of feedforward and feedback signals in the visual cortex to false percepts of oriented gratings were investigated using layer-specific fMRI in healthy human participants. If such false percepts are driven by feedback activity, they should be reflected by orientation-specific activity in the agranular layers of the visual cortex, whereas if false percepts are driven by feedforward activity they should be reflected by activity in the middle layers of the visual cortex. To preview, we replicate earlier findings that perceptual expectations are reflected by orientation-specific activity in the deep layers of the visual cortex^42^. However, these expectation signals did not affect the perceived orientation on high confidence false percept trials, suggesting these false percepts might not arise from feedback processes in the present paradigm. False percepts were instead reflected by orientation-specific activity in the middle layers. The involvement of the middle layers, taken together with a lack of expectation effects on perception in this experiment, suggests that false percepts can at least in principle arise from feedforward activity. This finding adds to the diversity of mechanisms that might underlie false perceptual inferences both in health and disease^43^.

## Results

### Experimental procedure

In the layer-specific fMRI study, 25 healthy participants were asked to complete a difficult perceptual discrimination task, in which they were required to report the orientation (45° or 135°) of a grating that was embedded in visual noise, as well as report their confidence that a grating was present (Fig. 1a). Unbeknownst to the participants, an auditory cue predicted the orientation of the upcoming gratings (Fig. 1b). On trials in which a grating was presented (grating-present trials, 50%), the auditory cues were 100% valid. Crucially, on the other 50% of trials, an auditory cue was followed by the presentation of visual noise only, and the gratings were omitted. Our research questions pertained to participants’ behaviour and how activity specific to perceived and expected orientations was represented in different cortical layers (Fig. 2) on these grating-absent trials. Specifically, we investigated whether participants reported high confidence false percepts on these trials, and if so, whether there was orientation-specific activity related to these false percepts in the feedback (agranular) or input (middle) layers of the early visual cortex.

**Fig. 1.**
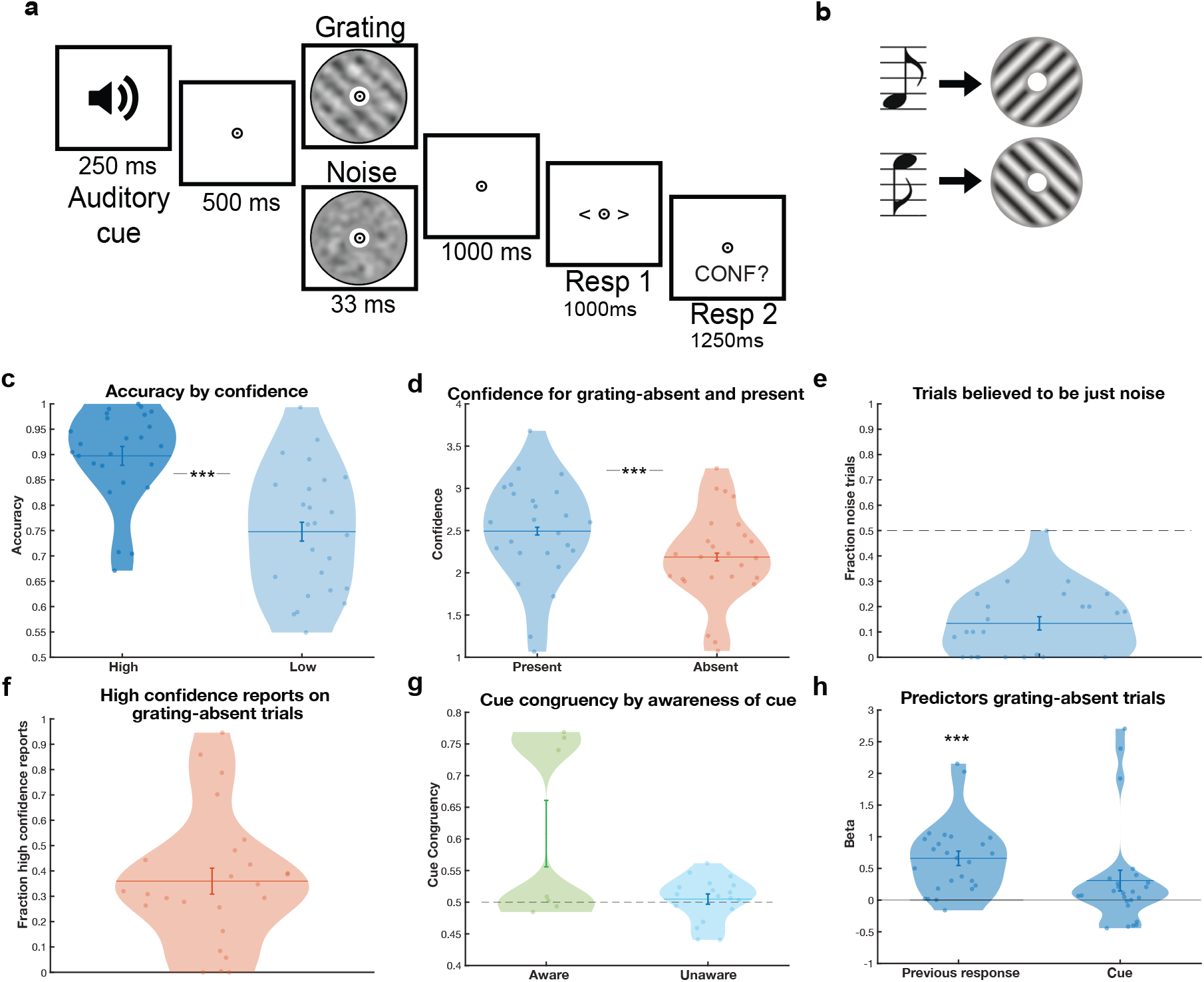
Experimental design and behavioural findings of layer-specific fMRI study. **a**, During the experiment an auditory cue was followed by either a low contrast grating embedded in noise (50% of trials), or a noise patch (50%). Participants indicated which orientation they saw and how confident they were that a grating was presented. **b**, One sound predicted the appearance of a 45 °, or clockwise, oriented grating, whilst the other predicted a 135 °, or anti-clockwise, orientated grating. Auditory cues were 100% valid on grating-present trials. **c**, On grating-present trials, participants’ orientation responses were more accurate when they indicated that they were confident (dark blue) compared to not confident (light blue) that they had seen a grating. **d**, Participants were more confident on grating-present trials (blue) compared to grating-absent trials (orange). **e**, Participants on average believed only ∼14% of trials to contain just noise (whereas the true proportion was 50%). **f**, On grating-absent trials, participants reported seeing gratings with high confidence (3 out of 4 or higher) on an average of 36% of trials. **g**, There was a slightly higher, non-significant, tendency to report orientations congruent with the expectation cue on grating-absent trials, which was driven by a few participants who were aware of the cue. **h**, Participants’ orientation response on the previous trial significantly predicted their orientation response on grating-absent trials. ***p<.001. Dots represent individual participants and violin shapes indicate density. Error bars indicate within-subject standard error of the mean for figure c & d, and standard error of the mean for e, f, g & h.

**Fig. 2.**
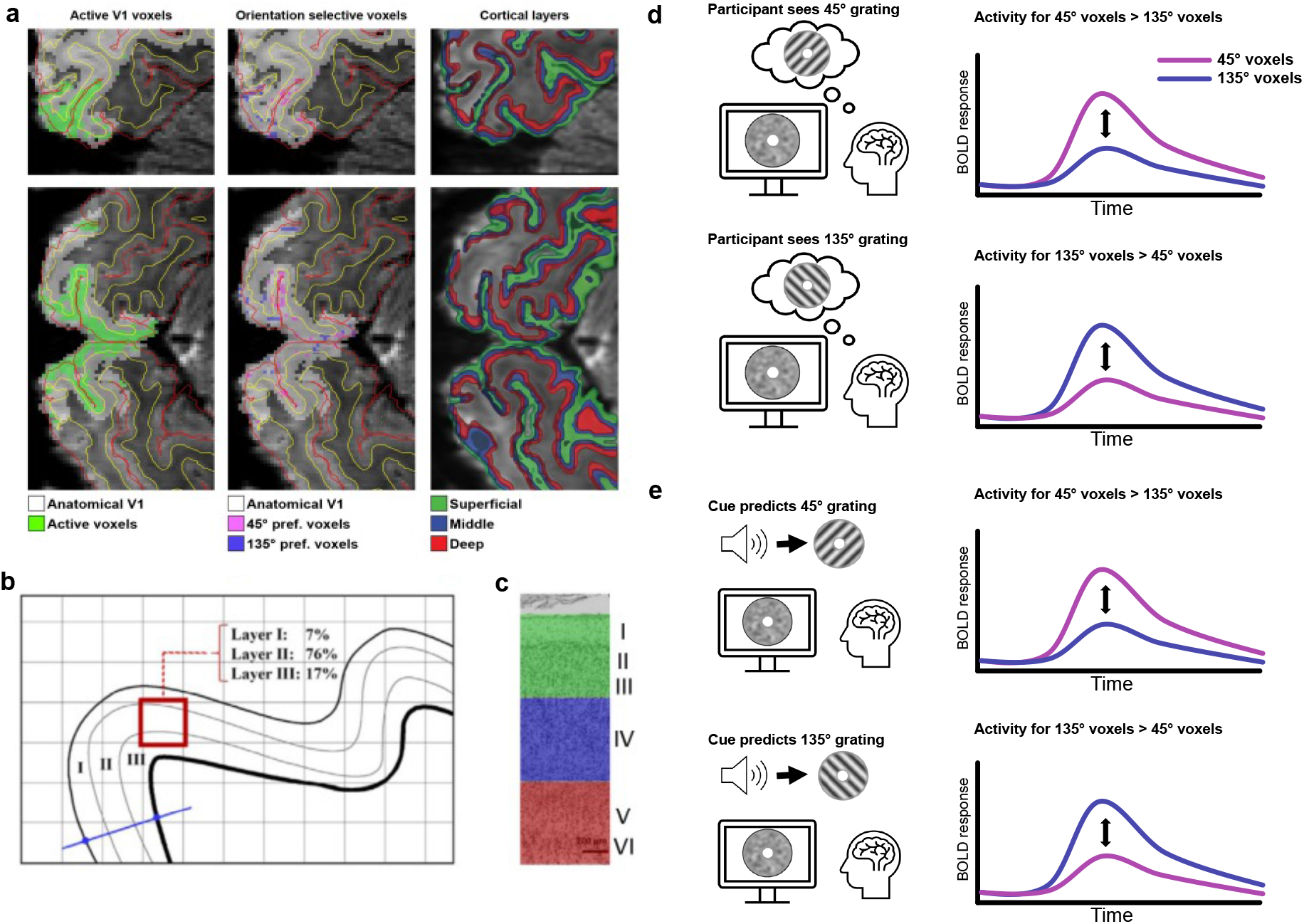
Schematic representation of analysis pipeline. **a (left)**, V1 and V2 voxels significantly activated by grating stimuli in a separate localiser run (green) were selected. **a (middle)**, Within this selection, the 500 voxels that mostly strongly preferred 45° over 135° gratings, as well as the 500 voxels that mostly strongly preferred 135° over 45°, were selected. **a (right), G**rey matter was divided into superficial, middle, and deep layers. **b**, For each of the 500 voxels for each orientation we calculated the contribution of each layer to that voxel. A spatial GLM was used to regress the contribution of each layer to the 500 voxels against the fMRI signal in these voxels, resulting in a single fMRI time course for superficial, middle, and deep layers for each set of orientation-preferring voxels. **c**, The superficial (green), middle (blue) and deep (red) layers roughly correspond to layers I to III, IV and V to VI respectively. Cytoarchitectural image of V1 adapted from ^84^ **d**, To calculate orientation-specific activity, BOLD activity was estimated for perceiving a 45° and 135° grating in both 45° and 135° preferring voxels, separately for each layer. Activity in the incongruent ROI (e.g., perceiving 135°, 45°-preferring voxels) was subtracted from congruent ROI effects (e.g., perceiving 45°, 45°-preferring voxels), which were then averaged over the two orientation-preference ROIs. This resulted in layer- and orientation-specific activity for perceived (d) orientations. **e**, The procedure was repeated for orientations predicted by the cue, resulting in layer and orientation specific activity for predicted orientations. Time courses on the right represent hypothesised BOLD responses to perceived and expected grating orientations in orientation-selective voxels.

### Participants experienced false percepts that were independent of perceptual expectation cues

Participants’ accurately identified the grating orientation on grating-present trials more often than expected by chance (mean accuracy =0.83, SD=.09) (T{24}=18.1, *p*<.001). Furthermore, they were more accurate when they were confident that they had seen a grating (i.e., higher than average confidence across trials) than when they were not (high: mean=0.90, SD=0.09; low: mean=0.75, SD=.12; paired t-test) (T{24}=5.7, *p*<.001) (Fig. 1c), demonstrating that they were able to perform the task and used the confidence ratings in a meaningful way. Participants also reported the perceived orientation more quickly on grating-present than grating-absent trials (F{1,24}=12.10, *p*=.002), as well as when they were more confident that they had seen a grating (F{1,24}=21.04, *p*<.001). Participants were more confident on grating-present trials (mean confidence =2.49, SD=.60, on a scale of 1-4) than grating-absent trials (mean=2.19, SD=0.54) (T{24}=4.76, *p*<.001) (Fig. 1d). Upon debriefing, all participants but one underestimated the frequency of grating-absent trials, believing on average that .14 (SD=.13) of trials contained just noise, while the true proportion was .50 (Fig. 1e). Strikingly, participants reported perceiving a grating with high confidence (3 out of 4 or higher) on 36% of grating-absent trials (Fig. 1f). Surprisingly, the perceptual expectation cues did not significantly bias which orientation participants perceived on grating-absent trials (0.53 false percepts congruent with the cue, chance level is 0.50, T{24}=1.90, *p* = 0.07). The small numerical trend towards false percepts being congruent with the expectation cues was driven by a few individuals who became aware of the meaning of the cues (N = 7 out of 25; Fig. 1g), potentially reflecting a concomitant response bias. High confidence false percepts were not more affected by the cues than low confidence, i.e. guessed, percepts (T{23}= 0.37, *p*=.71). Trial by trial predictors of participants’ choice behaviour were explored using a logistic regression model (see Methods for details). As expected, orientation responses on grating-present trials were predominantly driven by the presented stimulus (T{21}=12.6, *p*<.001), but also by which orientation was perceived on the previous trial, such that the previously reported stimulus was more likely to be perceived again on the current trial (T{21}=5.3, *p*<.001). Interestingly, on grating-absent trials, previous percepts also significantly predicted orientation reports (T{23}=5.48, *p*<.001), whereas the cues did not (T{23}=2.03, *p*=.056) (Fig. 1h). As above, the trend towards the cues influencing perception on grating-absent trials was driven by a few participants who became aware of the meaning of the cues. Thus, taken together, participants reported false percepts, but these were not significantly driven by the perceptual expectation cues in the present experiment, raising the possibility that these false percepts may have arisen from spontaneous fluctuations instead. We next investigated whether the false percepts were reflected in feedforward or feedback layers in the visual cortex.

### Estimation of layer- and orientation-specific activity

Ultra high-field (7T) fMRI was used to estimate orientation-specific activity in the deep, middle, and superficial layers of the early visual cortex (V1 and V2)^42,44,45^. To examine orientation-specific blood oxygenation level–dependent (BOLD) activity, V1 and V2 voxels were divided into two (45°-preferring and 135°-preferring) subpopulations based on an independent functional localiser. A cross-validation analyses confirmed that the grating stimuli presented in the localiser run induced stimulus-specific activity across all layers in V1 and V2 (All *p*<.05; see supplementary Fig. 1a). Layer-specific BOLD profiles were estimated for these subpopulations separately (Fig. 2). Importantly, we separately modelled the expected orientation of the grating, as defined by the cue, as well as the perceived orientation, defined by the participants responses, for each trial on grating-absent trials (see Methods for details). In other words, we were able to estimate the laminar profile of expected as well as perceived orientations using the same set of trials. Orientation-specific laminar BOLD profiles were then calculated by subtracting the laminar profile obtained from orientation-incongruent voxels (e.g., 135°-preferring voxels when a 45° grating was perceived) from the laminar profile in orientation-congruent voxels (45°-preferring voxels in this example). The resulting signal is a measure of how much more active neural populations in a particular layer were when their preferred grating orientation was perceived/expected, versus when their non-preferred orientation was perceived/expected. Crucially, this means that the layer-specific BOLD responses that are reported in the present study are specific to the perceived/expected grating orientation (as in^42,44^). In other words, BOLD responses reported for perceived (Fig. 3a) and expected (Fig. 3b) gratings reflect a neural representation of the specific orientation that was perceived/expected, rather than the overall, non-specific BOLD response.

**Fig. 3.**
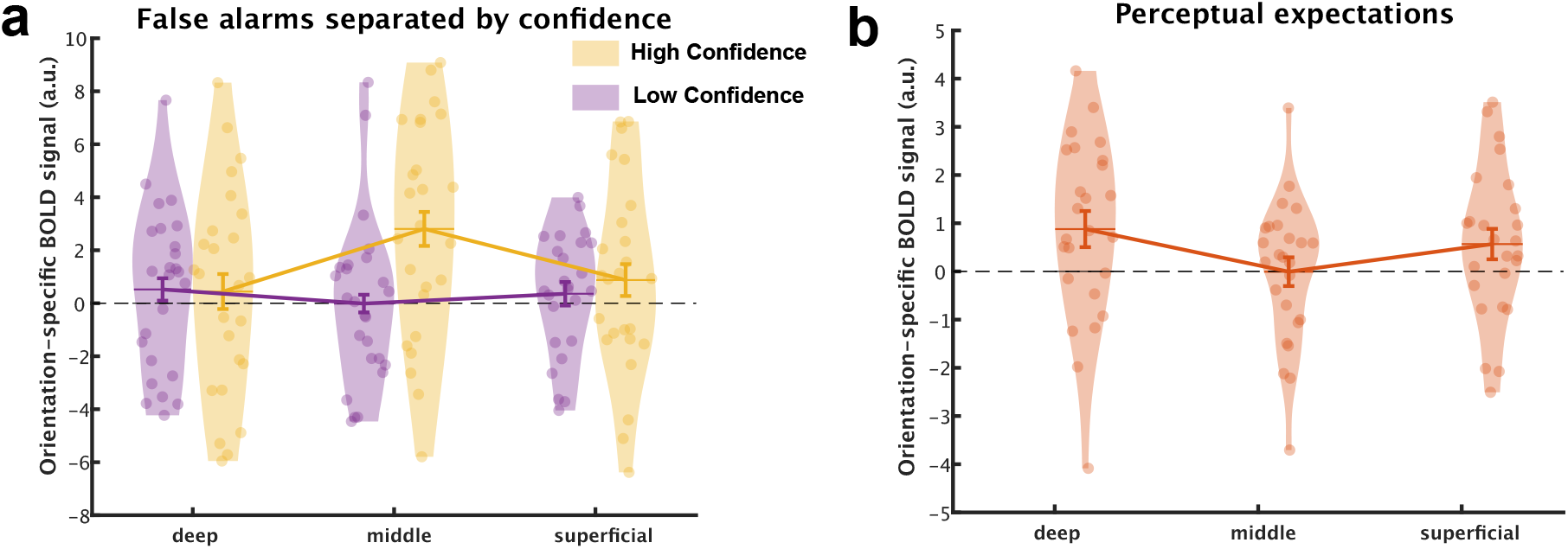
Orientation-specific BOLD activity in the cortical layers of V2. We calculated orientation-specific activity by separately estimating the effects of seeing or expecting 45° or 135° gratings, in voxels that preferred 45° or 135° gratings on the basis of an independent localiser, and subtracting incongruent activity (e.g. the effect of seeing 45° gratings in 135° preferring voxels) from congruent activity (the effect of seeing 45° gratings in 45° preferring voxels). Activity corresponding to perceived and expected gratings, respectively, were modelled separately on the same grating-absent trials. **a**, The middle layers contained orientation-specific activity reflecting high confidence false percepts (yellow), whilst low confidence trials (purple) did not induce orientation-specific activity in any of the layers. **b**, The deep and superficial layers reflected orientation-specific activity induced by perceptual expectations on grating-absent trials.Error bars represent within subject standard error of the mean.

### Orientation-specific activity reflecting false percepts and expectation cues in distinct cortical layers

In V2, the laminar profiles for false percepts with high confidence, false percepts with low confidence (guesses), and perceptual expectations, were significantly different from each other (interaction between three conditions and three layers: F{4,96}=4.39, *p*=.003; Fig. 3). High and low confidence false percepts were subsequently compared directly, revealing a significant difference in laminar profiles (interaction between two conditions and three layers: F{2,48}=5.42, *p*=.008). This was driven by increased orientation-specific activity for high compared to low confidence false percepts in the middle layers (paired t-test: T{24}=3.26, p=.003), but not in the superficial (T{24}=-0.73, *p*=.47) or deep (T{24}=-0.10, *p*=.92) layers (Fig. 3a). Thus, when participants reported perceiving a grating with a specific orientation with a high degree of confidence, even though no such stimulus was present, there was activity specific to the reported grating orientation in the middle layers of V2 (one-sample t-test: T{24}=3.43, *p*=.001). This high confidence false percept related activity in the middle layers was significantly higher than in the deep (T{24}=2.51, *p*=.019) and superficial layers (T{24}=2.24, *p*=.034). No orientation-specific activity was present when the percept was reported with low confidence, i.e., when the orientation report likely reflected a guess, rather than a genuine perceptual experience (one-sample t-test: T{24}=-0.21, *p*=.51). A direct comparison of high confidence false percepts and perceptual expectations demonstrated that they were associated with different laminar profiles (interaction between condition and layer F{2,48}=5.58, *p*=.007). This was primarily driven by a difference in middle layer activity (paired t-test: T{24}=3.34, *p*=.003), which were activated by high confidence false percepts (one-sample t-test: T{24}=3.43, *p*=.001) but not by expectations (one-sample t-test: T{24}=-0.19, *p*=.51). Conversely, perceptual expectations evoked significant orientation-specific activity in the deep layers (Fig. 3b) (one-sample t-test: T{24}=2.34, *p*=.014) and superficial layers (one-sample t-test: T{24}=1.8, *p*=.042). Conversely, false percepts did not activate deep (one-sample t-test: T{24}=0.56, *p*=.29) or superficial layers (one-sample t-test: T{24}=1.19, *p*=.12), although the difference between false percepts and expectation-induced activity was not significant in the superficial and deep layers (all *p>*.5).

In sum, high confidence false percepts were reflected by activity specific to the perceived orientation in the middle layers, whereas perceptual expectations were related to activity specific to the cued orientation in the deep layers. Interestingly, the effect of perceptual expectations in the deep layers was not driven by those participants who became aware of the meaning of the cues (two-sample t-test: T{23}=0.29, *p*=.78), and was significantly present in the subset of participants who were not aware of the cues’ meanings (N=18, T{17}=2.20, *p*=.021, see Supplementary Fig. 2c). Together, these findings suggests that false percepts can arise from feedforward activity within a cortical circuit different from the one signalling perceptual expectations.

The effects reported above for V2 did not extend to V1 (all *p*>.1). In fact, there was an interaction between layer (superficial, middle, and deep), stimulus condition (high and low confidence false percepts, and perceptual expectations), and ROI (V1 and V2) (F{4,96}=3.42, *p*=.012), in line with the effects being specific to V2. This is likely explained by the relatively low spatial frequency (0.5 cycles/°) gratings used here being more effective in activating V2 than V1, as has been demonstrated in animal studies^43^. In line with this, a cross-validated analysis of orientation-specific BOLD signals within the functional localiser revealed stronger orientation-specific effects in V2 than V1 across all layers (main effect of ROI: F{1,23}=23.56, *p*<.001, all layers *p*<.01); Supplementary Fig. 1a). Exploring the effects of the presented orientation on grating-present trials, we find that these low contrast gratings embedded in noise evoked significant orientation-specific activity in the superficial layers of V2 (T{24}=2.00, *p*=.028) but not the other layers (both *p*>.1; Supplementary Fig. 1b). It is perhaps not surprising that such weak and noisy stimuli did not evoke a significant orientation-specific BOLD signal in the deep and middle layers, but it is striking to note that more cognitive processes like expectation and perception did.

### Noise patch control analyses

Given that the false percepts on noise-only trials were reflected in the middle input layers of V2, one might be concerned that the noise patches themselves contained orientation signals that drove our effects. To avoid this issue a priori, we generated four noise patches with a flat orientation energy spectrum, as determined by a bank of orientation filters (based on Wyart et al. 2012; see Methods for details and Supplementary Fig. 2 for the stimuli used in the present study, and Supplementary Fig. 3 for the orientation energy profiles of the noise patches compared to the low-contrast gratings). However, participants were still significantly biased towards specific orientations for some of the noise patches. Specifically, noise patches two and three were significantly more often identified as 45° than 135° (63% and 69% respectively, T{24}=2.82, *p*=.01 & T{24}=5.3, *p*<.01), and vice versa for noise patch four (71%, T{24}=5.77, *p*<.01). Noise patch one showed no significant bias (54% identified as 135°, T{24}=1.31, *p*>.1). In theory, the stimulus-specific activity reported in the middle layers of V2 on the high confidence false alarm trials could therefore be driven by an unspecified signal present in the noise patches themselves. In order to investigate this possibility, we performed two control analyses.

First, if the noise patches were driving the results, the effects should disappear if all conditions of interest (i.e. low and high confidence false percepts of either orientation) were modelled separately for each noise patch, and then averaged over noise patches afterwards to ensure all patches contribute to both 45° and 135° percepts equally. In other words, in such an analysis each noise patch contributes equally to the estimated BOLD activity for 45° percepts and 135° percepts, regardless of participants’ propensity to perceive them as one more often than the other. In fact, this analysis replicated our core finding, namely that high confidence false percepts were reflected by increased middle layer activity compared to low confidence false percepts (T{24} = 2.51, *p*=.019). Indeed, there was significant orientation-specific activity in the middle layers for high confidence false percepts (T{24}=3.09, *p*=.003), but not in the other layers (p>.1) (see Supplementary Fig. 4). Therefore, when eliminating the possible confounding effect of the noise patches, our effects remain.

Second, we estimated any potential orientation-specific effects evoked by the different noise patches directly, comparing them against each other to explore any differences in stimulus-specific activity. We did not find any evidence that the noise patches alone caused stimulus-specific activity (across all combinations and layers *p*>.05; see Supplementary Fig. 5). Together, the results of these control analyses are not in line with an explanation of our effects in terms of stimulus-driven signals in the noise patches themselves, but instead strongly suggest that they reflect internally generated percepts.

### High confidence false percepts and reduced sensory precision predict everyday hallucination severity

In a separate online experiment (N=100), we tested whether the high confidence false percepts that were related to middle layer activity in the layer-specific fMRI study correlated with the prevalence and severity of hallucinatory percepts in daily life, as measured by the Cardiff Anomalous Perception Scale (CAPS) questionnaire^46^. The false percept task used here was similar to the one used in the fMRI experiment (with slight variations in practice procedure and trial counts; see Methods for details). One important difference was the introduction of three (rather than one) contrast levels on the grating-present trials, to enable estimates of sensory precision, i.e., how task accuracy depended on evidence quality. Specifically, a base-level contrast value was selected for each participant based on their performance during the practice phase, and this base-contrast was used during the main experiment along with gratings with 1% higher and 1% lower contrast.

Crucially, the prevalence of abnormal perceptual experiences in daily life (total CAPS scores^46^, was positively correlated with the average confidence that participants reported on grating-absent trials – i.e., the prevalence of high confidence false percepts in our task – across participants in the online sample (Rho=.22, *p*=.029 (Fig. 4a). Further, the sensory precision term – the influence of grating contrast on choice behaviour – correlated negatively with abnormal perceptual experience scores (Rho=-.30, *p*=.003) (Fig. 4b). In other words, the less sensitive participants were to stimulus contrast, the more likely they were to experience abnormal perceptual experiences in daily life. The effect of the expectation cues was not correlated with abnormal perceptual experience severity (*p*>.1). In general, the expectation cues had a very minor effect on choice behaviour, as in the fMRI experiment (see Supplementary Results; Supplementary Fig. 6). Using a linear regression model, both confidence on grating-absent trials (T{99}=2.01, *p*=.048) and sensory precision (T{99}=-2.98, *p*=.004) were found to be separate predictors of abnormal perceptual experience severity (overall linear regression model: F{2,97}=6.12, *p*=.003, R^2^=0.112). We did not find a relation between average confidence on grating-absent trials and delusion ideation (Rho=.13, *p*=.20), but there was a correlation with the sensory precision term (Rho=-.29, *p*=.004). In sum, high confidence false percepts of oriented gratings were related to the severity of everyday abnormal perceptual experiences. Given that the fMRI study showed that such high confidence false percepts were reflected by stimulus-like signals in the middle input layers of the early visual cortex, this raises the possibility that abnormal perception in everyday life may partly result from similar stimulus-like sensory fluctuations.

**Fig. 4.**
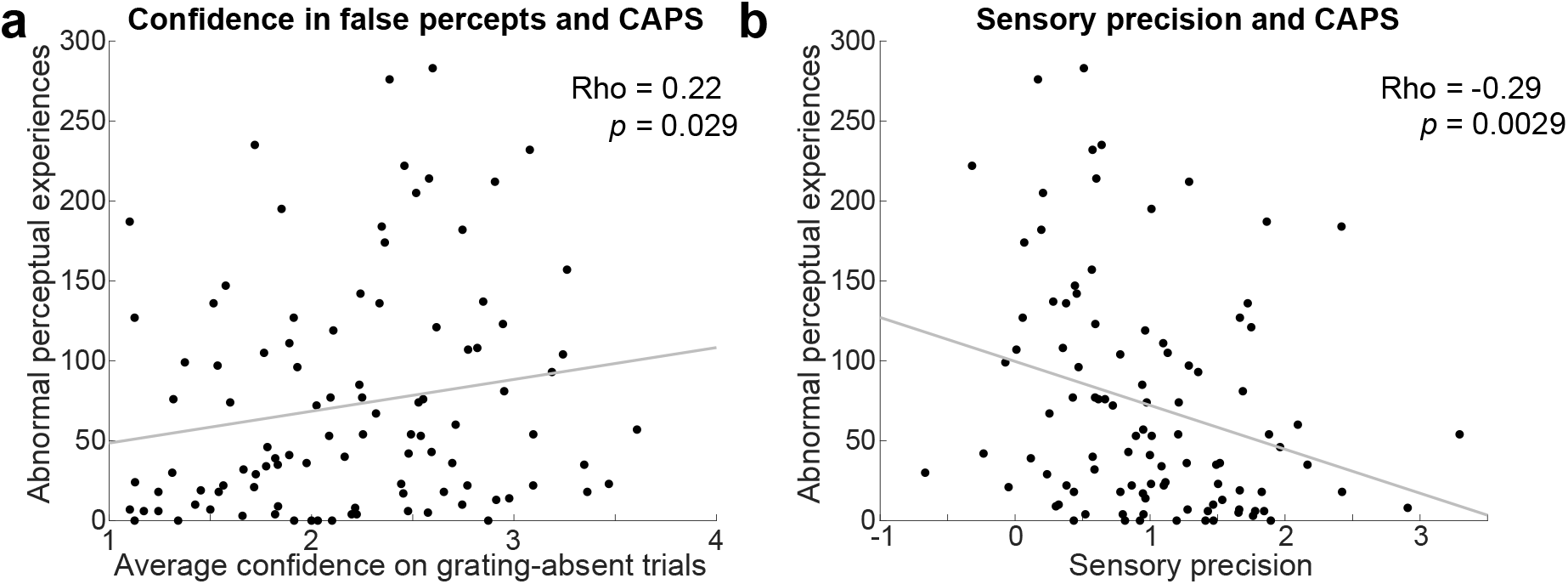
Correlations between grating percepts in the current task and everyday hallucinations. **a**, Abnormal perceptual experiences correlated with confidence in false percepts. **b**, Abnormal perceptual experiences correlated with less reliance on sensory precision (i.e., the interactive effect of grating contrast and orientation on choice behaviour).

## Discussion

There is a wide range of theories that attempt to explain the neural mechanisms of false percepts. The dominant theory highlights the role of feedback prediction signals in driving false percept ^2–5^. Others have proposed that false percepts can emerge in feedforward fashion as well^19,20^. Here, we tested feedforward and feedback contributions to false percepts using layer-specific fMRI. We found that the deep layers of the visual cortex reflected perceptual expectations, replicating previous research^42^. However, high confidence false percepts, were reflected solely in the middle input layers of the early visual cortex, and were unaffected by perceptual expectation cues. These high confidence false percepts correlated with everyday hallucination severity in a separate online study. This suggests that false percepts can arise from stimulus-like feedforward activity in sensory cortex, and do not necessarily require feedback-induced stimulus templates in the deep layers of the visual cortex.

The core finding here – that orientation-specific activity in the middle layers of V2 can lead participants to perceive a grating that was not actually presented – has important implications for the field of hallucination research as well as perception research more generally. That is, it suggests that false percepts can arise through activity in the input layers, in the absence of top-down stimulus templates induced by perceptual expectations. This puts a larger emphasis on feedforward signals than previously assumed in dominant models of hallucinations, which have largely focused on the role of top-down expectations in driving false percepts^2,3,5,8,10,11,17^. Thus, this research adds to the diversity of mechanisms that might underlie false inferences. These findings are in line with theories that state that false percepts can arise from spontaneous activity that resembles sensory input^20^, which recent studies have confirmed in neurological disorders like Charles Bonnet Syndrome^19^. Furthermore, they expand upon previous studies that have found that early sensory activity can lead to false alarms^28–32,47^, by suggesting that the activity reported by these studies may reflect spontaneous fluctuations in the input layers, rather than feedback from higher-order regions. Recent circular inference models of hallucinations have emphasised the role of ascending loops, akin to feedforward activity, in unimodal hallucinations as seen in psychotic disorders^23,48^. Specifically, they suggest that weak sensory signals can trigger perceptual hypotheses that are then counted as sensory evidence themselves in runaway ‘overcounting’ loops. This overcounting of sensory signals has been shown to correlate with positive symptoms (e.g., hallucinations and delusions) in schizophrenia patients^24^.

Top-down processes like visual working memory and perceptual expectations induce stimulus representations in the deep and superficial, but not the middle layers of the early visual cortex^42,45,49^. However, keeping an image in visual working memory or merely expecting a visual stimulus does not lead to a concurrent perceptual experience, as is the case with a hallucination. One interpretation therefore might be that the middle layers are required to be activated in order for a perceptual experience to occur that is attributed externally. Indeed, it has been suggested that feedforward signals are essential in distinguishing imagination from veridical perception^50^. Therefore, even hallucinations that are induced by top-down perceptual expectations might do so through their modulation of middle layer activity. Further research is needed to answer this question.

The prevalence of high confidence false percepts in our experiment correlated with everyday abnormal perceptual experiences. This is in line with previous studies demonstrating that those who hallucinate are more prone to perceive stimuli in noise in detection tasks^10,11,13,18^. Furthermore, higher hallucination scores correlated negatively with the sensory precision term of our logistic regression model. That is, the less participants relied on the contrast of the sensory stimulus in making their perceptual decision, the more they experienced hallucinations in everyday life. This is in line with Bayesian models of hallucinations, where a reduction in sensory precision increases the influence of prior expectations, possibly leading to hallucinations^51^. Interestingly, reduced reliance on sensory contrast on the one hand, and confidence on grating-absent trials on the other, were separate predictors of everyday hallucinations severity, suggesting separate underlying mechanisms contributing to hallucinations. The present study revealed that activity fluctuations in the middle input layers reflected high confidence false percepts. The strength of such fluctuations in an individual, in combination with a loss of precision in sensory encoding in these same input layers, might particularly amplify the tendency to experience hallucinations in everyday life. This is in line with the idea that a loss of sensory precision is a separate compounding factor, which allows hallucinations to manifest^52^. In psychiatric conditions, this might come in the form of internal noise in the sensory systems. In neurological disorders, this is most prominently seen in Charles Bonnet syndrome^20,53^.

These findings should not be taken as evidence against the theory that top-down perceptual expectations can play an important part in generating hallucinations, for which there is ample indirect evidence^10–14,17,18,54^. Instead, these findings highlight that the possibility that false percepts could occur in principle from feedforward activity in early sensory regions. Higher-order cortical areas are likely to be involved in interpreting this feedforward stimulus-specific activity.

We speculate that there might be two different types of hallucinations: feedforward driven hallucinations, and hallucinations driven by top-down perceptual expectations. These might reflect phenomenologically different hallucinations, potentially mapping onto so-called minor phenomena and complex visual hallucination respectively^55,56^. There is some evidence that these reflect different neural processes in Lewy body dementia^57^. The hallucinations reported in the current study may be more akin to minor phenomena, reflecting low-level orientation-specific activity resulting in low-level visual distortions. Those driven by perceptual expectations could result in more complex visual hallucinations, and might in contrast be reflected by deep layer activity. Future studies should aim to study the laminar profile of hallucinations elicited by perceptual expectations to test these hypotheses.

An alternative possibility is that top-down perceptual expectations might still play a role in the present study through the experimental set-up. Specifically, on every trial there was likely a strong expectation that one of two stimuli would be presented. This is underlined by the debriefing questionnaire, where participants greatly underestimated the proportion of trials on which the gratings were absent, suggesting that they expected to see gratings on most trials, regardless of their specific orientation. This expectation about stimulus presence versus absence could be an important driver of false percepts. Indeed, it is typically the expectation of the likelihood of the presence of a stimulus, rather than its specific contents, that has been found to be associated with psychosis in previous studies^10,11,17^. Interestingly, expectations about stimulus presence versus absence and expectations about stimulus content have been suggested to be supported by different neural processes^32,58,59^. Future research could elucidate the contribution of stimulus presence expectations on the effects found in the present study by manipulating this explicitly.

In theory, the V2 middle layer signals reflecting false percepts in the current study could have resulted from feedback signals to V1 or the thalamus being sent downstream to the middle layers of V2. Such an indirect feedback effect does not seem in line with the absence of stimulus-specific effects reflecting false percepts in V1, but it should be acknowledged that indirect feedback effects cannot be fully ruled out on the basis of a null effect.

On a neural level, an expectation of stimulus presence might prime pyramidal neurons in the deep cortical layers through receptors on their apical dendrites^60,61^. Targeting the apical dendrites is expected not to drive these pyramidal neurons directly, but allow them to function as coincidence detectors^61^. In turn, deep layer neurons can modulate incoming sensory input through their projections onto the middle layers^62,63^. This allows sensory input concurrent with expectations to be processed more quickly, giving them a head start in signal processing^63–65^. Furthermore, once evidence accumulates for one orientation in the middle layers, possibly due to random fluctuations, the evidence for the other orientation may be suppressed through lateral inhibition^66,67^. This hypothesised circuit has the potential to implement circular inference, whereby top-down expectations can get counted as sensory evidence in ascending loops^23,24,68^. This would subsequently result in orientation-specific activity in the middle layers driving hallucinations as seen in our study. This modulatory (rather than driving) role of feedback connections^63^ may explain why many studies have found that concurrent noisy input is required to induce hallucinations^10,11,13–15,17,18^. Alternatively, the prior on stimulus presence may not have affected sensory cortex itself, but instead biased source monitoring processes in the prefrontal cortex^50^ towards judging spontaneous fluctuations in the middle layers of V2 as being externally generated.

Why did the expectation cues not induce false percepts in the current study? We suggest this may be due to recruiting a normative sample who might rely less on expectations than those prone to hallucinate^10,13,15^. Second, there is increasing evidence that conscious expectations exert stronger effects on perception than unconscious expectations^69,70^. Since for the majority of individuals in this experiment the expectations were implicit, this might have weakened their effect. Studies that do report effects of implicit cues on perception typically reveal biased perception of existing stimuli, rather than eliciting percepts de novo^42,71^ although see^72^ for a notable exception.

Despite the expectation cues not affecting perception, they did induce orientation-specific templates in the deep layers of the early visual cortex, in line with previous work^42^. Interestingly, in contrast to the previous study, there was also significant expectation-evoked activity in the superficial layers. This additional effect might be explained due to the concurrent presentation of noisy stimuli in this study, where no stimulus was presented in the previous study. That is, speculatively, the presence of (noisy) sensory input may ‘unlock’ modulatory effects of feedback in the superficial layers. Strikingly, the representation of expected orientations in the deep layers was reliable even in those who were not aware of the cue-stimulus relationship, which suggests that the brain can generate sensory expectations based on statistical relationships that are learnt outside of conscious awareness^73^. The finding that orientation-specific deep layer activity might not be sufficient to generate a conscious percept seems in conflict with theories that emphasise the importance of deep layer activity in generating conscious percepts^74,75^. Our work nuances that picture, by suggesting that deep layer activity by itself may not be sufficient for conscious awareness. Of course, we cannot rule out that the deep layer activity revealed here was simply not strong enough to induce conscious percepts.

In the present study, no reliable effects of either perceptual expectations or false percepts were found in V1. This is likely due to the lower spatial frequency stimuli used in the present study (0.5 cycles/°) compared to previous studies that reported orientation-specific effects in V1 (1.0-1.5 cycles/°;^44,45,76^), as V2 neurons prefer lower spatial frequencies than V1 neurons^77^. This was confirmed by our localiser analyses, showing stronger orientation-specific effects in V2 than in V1. While the localiser task did induce orientation-specific effects in V2, no stimulus-driven effects were found for the presented stimuli in the main task, likely due to these being very low contrast and embedded in noise. This suggests that neural signals induced by expectations and falsely perceived gratings were more reliable than those evoked by very noisy bottom-up inputs, in line with a previous study that also found orientation-specific activity reflecting false percepts, but not presented stimuli embedded in noise^31^.

In conclusion, high confidence false percepts were reflected by orientation-specific activity in the middle input layers of the early visual cortex, whereas perceptual expectations activated the deep layers. These findings suggest that false percepts can arise from low-level content-specific fluctuations in the input layers of the visual cortex. This nuances the view that false percepts are necessarily driven by top-down expectations^2,3,5^. Future studies should aim to further explore the nature of these low-level fluctuations and what drives them, as well as investigate whether false percepts can also be driven by purely top-down signals. These findings have important implications for our understanding of the neural mechanisms underlying hallucinations, revealing how the brain can generate perception in the absence of sensory input.

## Acknowledgements

The authors would like to thank Fraser Aitken for help with early piloting of the experimental paradigm. This work was supported by a Wellcome/Royal Society Sir Henry Dale Fellowship [218535/Z/19/Z] and a European Research Council (ERC) Starting Grant [948548] to P.K. The Wellcome Centre for Human Neuroimaging is supported by core funding from the Wellcome Trust [203147/Z/16/Z].

## Competing interests

The authors declare no competing interests

## Methods

### Ethics statement

This study was approved by the University College London Research Ethics Committee (R13061/RE002 for the imaging study, R6649/RE004 for the online study) and was conducted according to the principles of the Declaration of Helsinki. All Participants gave written informed consent prior to participation and received monetary compensation (£7.50 an hour for behavioural tasks, £10 an hour for MRI)

### Participants

Twenty-eight healthy human volunteers with normal or corrected-to-normal vision participated in the 7T fMRI experiment. Three participants were excluded due to our strict head motion criteria of no more than 10 movements larger than 1.0 mm in any direction between successive functional volumes. For the remaining participants, the maximum change in head position in any direction over the course of the fMRI runs was within 4 mm (0.66 +/− 0.54 mm, mean +/− SD over participants) of the mean head position (to which the anatomical boundaries were registered). The final sample consisted of 25 participants (22 female; age 25 ± 4 years; mean ± SD).

One hundred participants participated in the online study. Participants were recruited through Prolific (www.prolific.co) and were paid £7.50 for their participation. Three participants were excluded for failing to answer the catch questions on the questionnaire correctly, resulting in a final sample of 97 participants.

### Questionnaires

For the online study questionnaire data was collected for the Peter Delusions Index^78^, as well as the Cardiff Abnormal Perception Scale^46^. Total scores were calculated for the PDI and CAPS by adding their respective subscales. These were then correlated with the behavioural measures for the online study.

### Stimuli

Grayscale luminance-defined sinusoidal Gabor grating stimuli were generated using MATLAB (MathWorks, Natick, Massachusetts, United States of America, RRID:SCR_001622) and the Psychophysics Toolbox (Brainard, 1997). During the behavioural session for the fMRI study, the stimuli were presented on a PC (1920 × 1200 screen resolution, 60-Hz refresh rate). In the fMRI scanning session, stimuli were projected onto a rear projection screen using an Epson EB-L1100U Laser projector (1920 × 1200 screen resolution, 60-Hz refresh rate) and viewed via a mirror (viewing distance 91 cm). On grating-present trials (50%), auditory cues were followed by a grating (0.5-cpd spatial frequency, 33-ms duration, and separated by a 750-ms blank screen), displayed in an annulus (outer diameter: 10° of visual angle, inner diameter: 1°, contrast decreasing linearly to 0 over 0.7° at the inner and outer edges), surrounding a fixation bull’s eye (0.7° diameter). These stimuli were combined with one of 4 noise patches, which resulted in a 4% contrast grating embedded in 20% contrast noise during the fMRI session. On grating-absent trials, one of the 4 noise patches was presented on its own. Noise patches were created through smoothing pixel-by-pixel Gaussian noise with a Gaussian smoothing filter, ensuring that the spatial frequency of the noise patches matched that of the gratings. This was done to ensure that the noise patches and gratings had similar low-level properties, increasing the likelihood of reporting false percepts. To avoid including noise patches which contained grating-like orientation signals by chance, the noise patches were processed through a number of Gabor energy filters with varying preferred orientations. The noise patches with low (2%) signal energy were selected to be included for the present experiment (see supplementary Fig. 3 & Fig. 4). The resulting four noise patches were used for all participants throughout the experiment, ensuring that reported false percepts could only be triggered by internal mechanisms^31,79^. During the practice session on the first day, the contrast of the gratings was initially high (80%), gradually decreasing to 4% towards the end of the practice. The central fixation bull’s-eye was present throughout the trial, as well as during the intertrial interval (ITI; jittered exponentially between 2,150 and 5,150 ms). In the online study, multiple grating contrast levels were presented, titrated to each individual (see procedure below).

### Experimental procedure

On the first day of testing for the laminar fMRI study, participants underwent a behavioural practice session. The practice consisted of an instruction phase with 7 blocks of 16 trials where the task was made progressively more difficult, whilst verbal and written instructions were provided. During the practice runs, the auditory cues predicted the orientation of the first grating stimulus of the pair with 100% validity (45° or 135°; no grating-absent trials). After the completion of the instructions, the participants completed 4 runs of 128 trials each, separated into 2 blocks of 64 trials each. In the first 2 runs the expectation cues were 100% valid, to ensure participants learnt the association, whilst in the final 2 runs the cue was 75% valid, to test whether participants might have adopted a response bias. Grating contrast decreased over the 4 runs, specifically the contrast levels were 7.5, 6, 5, and 4%, while the contrast of the noise patches remained constant at 20%. No grating-absent trials were presented on day 1. On the second day, participants performed the same task in the MR scanner. As on the first day, 4 runs were completed, but now the grating contrast was fixed at 4% on grating-present trials, and on 50% of the trials the gratings were omitted and only noise patches were presented, resulting in grating-absent trials. Each run lasted ∼12.5 minutes, totalling ∼50 minutes.

Trials consisted of an auditory expectation cue, followed by a grating stimulus embedded in noise on 50% of trials (750-ms stimulus onset asynchrony (SOA) between cue and grating). The auditory cue (high or a low tone) predicted the orientation of the grating stimulus (45° or 135°). On grating-present trials, a grating with the orientation predicted by the auditory cue was presented embedded in noise, while on grating-absent trials only a noise patch was presented. The stimulus was presented for 33ms in the fMRI study. After the stimulus disappeared, the orientation response prompt appeared, consisting of a left and right pointing arrow on either side of the fixation dot (location was counterbalanced). Participants were required to select the arrow corresponding to their answer (left arrow for anticlockwise, or 135°, right arrow for clockwise, or 45°; 1s response window) through a button press with their right hand. Subsequently the letters “CONF?” appeared on the screen probing participants to indicate their confidence that they had seen a grating (1 = I did not see a grating, 2 = I may have seen a grating, 3 = I probably saw a grating, 4 = I am sure I saw a grating), using 1 of 4 buttons with their left hand (1.25s response window). Participants indicated their response using an MR-compatible button box in the MRI scanner, and a keyboard during training.

After the main experiment, participants performed a functional localiser task inside the scanner. This consisted of flickering gratings (2 Hz), presented at 100% contrast, in blocks of approximately 14.3 seconds (4 TRs). Each block contained gratings with a fixed orientation (45° or 135°). The 2 orientations were presented in a pseudorandom order followed by an approximately 14.3-second blank screen, containing only a fixation bull’s-eye. Participants were tasked with responding whenever the black fixation dot briefly dimmed to ensure central fixation. All participants were presented with 16 localiser blocks which totalled approximately 15 minutes.

The online study was created and hosted using Gorilla Experiment Builder (www.gorilla.sc)^80^. Participants were recruited through Prolific (www.prolific.co). Prior to the start of the instruction blocks participants were asked to keep a 50cm distance from the screen. They were required to adjust the size of a rectangle on their screen to match a bank card, to ensure that the visual angle was equal across participants. Subsequently, they were asked to adjust the volume of their headphones to a high but not unpleasant volume. The instruction phase of the online study was the same as the instruction phase of the fMRI study. The timings for the trials were identical as well, except that the stimuli appeared on the screen for 50ms instead of 33ms, due to software constraints. After completing the 7 instruction blocks, participants were required to complete 4 blocks with different grating contrast levels (7.5, 6, 5, and 4%, in that order). The lowest grating contrast for which participants were able to perform the orientation task with at least 75% accuracy served as the base contrast value for the main experiment. During the main experiment, participants were required to complete 4 blocks of 128 trials. Unlike in the fMRI experiment, different grating contrast levels were presented, and expectation cues were sometimes invalid (6.7% of grating present trials). Specifically, out of the 128 trials on each block, on 96 trials (75%) a grating was present and on 32 (25%) only a noise patch was presented. Of the 96 grating-present trials, one third (32) were presented at the base contrast, one third at base - 1% contrast, and the other third at base + 1% contrast. Of these grating-present trials, 90 (93.3 %) were valid and 6 (6.7 %) were invalid.

### fMRI data acquisition

MRI data were acquired on a Siemens Magnetom Terra 7T MRI system (Siemens Healthcare GmbH, Erlangen, Germany) with an 8 channel head coil for localised transmission, operating in a quadrature-like (‘TrueForm’) mode, with a 32-channel head coil insert for reception (Nova Medical, Wilmington, USA) at the Wellcome Centre for Human Neuroimaging (University College London). Functional images were acquired using a T2*-weighted 3D gradient-echo EPI sequence (volume acquisition time of 3,552 ms, TR = 74 ms, TE = 26.95 ms, voxel size 0.8 × 0.8 × 0.8 mm^3^, 15° flip angle, field of view 192 × 192 × 38.4 mm^3^, GeneRalized Autocalibrating Partial Parallel Acquisition (GRAPPA) acceleration factor 4, and partial Fourier 6/8 in the phase-encoded direction of the EPI readout, binomial (1331) water-selective excitation). Anatomical images were acquired using a Magnetization Prepared 2 Rapid Acquisition Gradient Echo (MP2RAGE) sequence (TR = 5,000 ms, TE = 2.54 ms, TI = 900 ms and 2,750 ms, voxel size 0.65 × 0.65 × 0.65 mm, 5° and 3° flip angles, field of view 208 × 208 × 156 mm^3^, in-plane GRAPPA acceleration factor 3).

### Preprocessing of fMRI data

*T*he first 2 volumes of each run were discarded. Prior to registration the functional volumes were cropped to cover only the occipital lobe to reduce the influence of severe distortions in the frontal lobe. The cropped functional volumes were spatially realigned within scanner runs, and subsequently between runs, to correct for head movement, using SPM12 (https://www.fil.ion.ucl.ac.uk/spm/).

### Segmentation and coregistration of cortical surfaces

The methods for segmenting and coregistering cortical surfaces are identical to several previously published studies^36,42,44,45^, and are reiterated here. Freesurfer (http://surfer.nmr.mgh.harvard.edu/) was used to detect the grey (GM) and white matter (WM) boundaries and cerebral spinal fluid (CSF) based on a bias corrected MP2RAGE image. The boundaries were checked for errors where the dura was mistakenly included in the pial surface. Subsequently the GM boundaries were registered to the mean functional image. Specifically, a conventional rigid-body registration was followed by a recursive boundary-based registration (RBR)^81^. RBR consisted of applying boundary-based registration (BBR) recursively to increasingly smaller partitions of the cortical mesh. An affine BBR was applied with 7 degrees of freedom: rotation and translation along all 3 dimensions and scaling along the phase-encoding direction only. This scaling allows correction of distortions along the low bandwidth phase-encoded EPI direction of acquisition. In each iteration, the cortical mesh was split into 2, and the optimal BBR transformations were found and applied to the respective parts. Subsequently, each part was split into 2 again and registered. The specificity increased at each stage and corrected for local mismatches between the structural and the functional volumes that are due to magnetic field inhomogeneity-related distortions. Six such iterations were performed. The splits were made along the cardinal axes of the volume, such that the number of vertices was equal for both parts. The plane for the second cut was orthogonal to the first, the third was orthogonal to the first 2. The median displacement was taken after running the recursive algorithm 6 times, in which different splitting orders where used, comprised of all 6 permutations of x, y, and z.

### Definition of regions of interest

The definition of regions of interests (ROIS) was identical to a recently published study^42^, and reiterated here. V1 and V2 surface labels were obtained through Freesurfer, based on the segmentation of the MP2RAGE image. These were subsequently projected to volume space, covering the full cortical depth plus a 50% extension into WM and CSF. The V1 and V2 ROIs were subsequently constrained to the voxels that were responsive to the localiser gratings. Specifically, separate regressors were defined for the blocks of 45° and 135° gratings, respectively, and the mean of the resulting parameter estimates was contrasted against baseline to identify voxels that exhibited a significant response to the grating stimuli irrespective of orientation (T > 2.3, *p* < 0.05; V1: mean=6208, SD=1799 voxels; V2: mean=9370, SD=3256 over participants). Subsequently, the orientation preference of each voxel was estimated by contrasting the 2 orientation regressors. The 500 voxels that most strongly favoured the 45° and 135° gratings, respectively, constituted the two orientation-specific ROIs within V1 and V2. Finally, each voxel’s time course was normalised (z-scored), and multiplied by the absolute T-value of the orientation contrast (45° versus 135°), to weight the data by the most robust orientation preference. Note that all reported effects in the z-scored data were also present without z-scoring. These ROI definitions were identical to those used in previous studies that successfully resolved orientation-specific BOLD signals with layer specificity ^42,44,45^. The analysis approach was matched to these previous studies to facilitate comparisons between previous findings that involve orientation- and layer-specific fMRI signals.

### Definition of the cortical layers

GM was divided into 3 equivolume layers using the level set method described in detail elsewhere^81–83^, following the principle that the layers of the cortex maintain their volume ratio throughout the curves of the gyri and sulci^83^. Briefly, the level set function is a signed distance function (SDF), where points on the same surface equal 0 and values on one side of the surface are negative and values on the other are positive. The level set function for the GM–CSF and GM–WM boundaries is calculated, and then intermediate surfaces can be defined by moving the surface to intermediate cortical depths. The equivolume model transforms a desired volume fraction into a distance fraction, taking the local curvature of the pial and WM surfaces at each voxel into account^81^. Two intermediate surfaces between the WM and pial boundaries were calculated, yielding 3 GM layers (deep, middle, and superficial). In human early visual cortex, these 3 laminar compartments are expected to correspond roughly to layers I to III, layer IV, and layers V and VI, respectively^84^. Based on these surfaces, 4 SDFs were calculated, containing for each functional voxel its distance to the boundaries between the 5 compartments (WM, CSF, and the 3 GM layers). This set of SDFs (or “level set”) allowed the calculation of the distribution of each voxel’s volume over the 5 compartments^81^. This layer volume distribution provided the basis for the laminar GLM discussed below.

### Extraction of layer-specific time courses

Because the fMRI data consisted of 0.80 mm isotropic voxels, individual voxels will naturally contain signals from multiple layers, as well as WM and CSF. Thus, if we were to simply interpolate the fMRI signal at different depths, there will be contamination from bordering layers. One way to address this so-called partial volume problem is to decompose the layers by means of a spatial GLM^36,37,42,81^. For every ROI, a laminar design matrix **X** represents the distribution of the 500 voxels over the different layers (n x k, where n = 500 voxels, and k = 5 laminar compartments). Every row of **X** indicates the proportions of the layers covered by a particular voxel, and the columns represent the volume of the corresponding layer across voxels. This laminar design matrix can be used in a spatial GLM to separate the BOLD signal of the 5 different laminar compartments (three GM layers, WM, and CSF) through ordinary least squares (OLS) regression^81^:

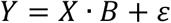

Here **Y** is a vector of voxel values from an ROI in a specific functional volume, **X** is the laminar design matrix, and **B** is a vector of layer signals. For each ROI and each functional volume, the layer signal 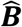 was estimated by regressing **Y** against **X**, yielding 5 depth-specific time courses per ROI.

To confirm that the method correctly identified GM, the raw signal in the EPI volumes for each of the 3 GM layers was quantified, as well as WM and CSF. As expected, the signal intensity was higher in the 3 GM layers (deep: 239 +/− 38; middle: 240 +/− 46; superficial: 239 +/− 40; mean +/− SD over participants) than in WM (209 +/− 30) and CSF (230 +/− 51) (T{24}= 5.69, *p* = 7.4 × 10^−6^).

### Estimating effects of interest per layer

A temporal GLM was used to estimate the effects of interest in each of the 3 GM layers. A model with 4 regressors of interest was used to estimate the effects of perceptual expectation (grating-present and grating-absent trials, separately for the 45° or 135° orientations). These regressors of interest were constructed by convolving stick functions representing the onsets of the trials with SPM12’s canonical haemodynamic response function as well as their temporal derivative, resulting in Beta values for each experimental effect. Furthermore, the head motion parameters, their derivatives, and the square of the derivatives, were included as nuisance regressors. Subsequently, the data and the design matrix were high-pass filtered (cut-off = 128 seconds) to remove any low-frequency signal drifts.

In order to calculate orientation-specific BOLD responses, the layer-specific parameter estimates for each orientation in the non-corresponding ROI (e.g., a 45° grating/expectation in a 135°-preferring ROI) were subtracting from the parameter estimates in their corresponding ROI (e.g., a 45° grating/expectation in a 45°-preferring ROI; see equation below where B stands for Beta).

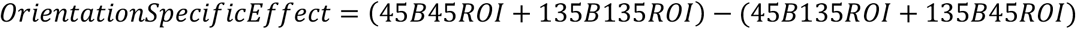

This procedure was followed for all the laminar analyses presented in this study (perceptual expectations, high confidence false percepts, low confidence false percepts). These estimated BOLD responses were subjected to a 2-way repeated measures ANOVA with factors perceptual condition (expectation, high confidence false percept, low confidence false percept), and cortical layer (deep, middle, superficial). The main effect of interest, namely whether laminar BOLD profiles differed for perceptual expectations and hallucinated gratings, was tested by the interaction of perceptual condition and cortical layer. To follow up a significant effect with all 3 perceptual conditions included, further repeated measures effects were performed to specifically test 1) the interaction between perceptual expectation vs high confidence false percept and cortical layer to explore whether hallucinations were specifically different from perceptual expectations, and 2) the interaction between high and low confidence false percepts and cortical layer to explore whether being confident in a false percept affects the laminar profile. Significant interactions were followed up with paired-sample *t* tests. Finally, orientation-specific effects in specific layers were tested against zero using one-sample t-tests (one-tailed). To visualise the relevant across-subject variance for the within-subject ANOVA, errors bars in all figures show within-subject standard error of the mean (SEM)^85,86^.

### Behavioural analyses

For the online study, accuracy and confidence scores were compared across the different contrast levels using repeated measures ANOVAs. Accuracy was also compared between the different confidence levels, to test whether participants were more accurate at identifying grating orientation when they were more confident that they had seen a grating. The effect of the expectation cues was assessed by exploring whether participants tended to report orientations in line with the cue. Follow-up tests were performed to investigate whether the cues’ effects were mediated by awareness of their meaning. To understand what drives abnormal perceptual experiences, a logistic regression model was used to explore which factors predicted orientation responses on grating-present and grating-absent trials separately. Predictors for grating-present trials were current stimulus orientation, current stimulus contrast, orientation predicted by the cue, orientation response on the previous trial, and the interaction between present stimulus contrast and orientation (as a measure of sensory precision). For the grating-absent trials, the predictors included previous orientation response, and orientation predicted by the cue (as there was no present stimulus orientation or contrast). Finally, we tested whether abnormal perceptual experiences as measured using the CAPS questionnaire were correlated with cue effects, confidence on grating-absent trials, and sensory precision, using Spearman’s rank correlation.

Similarly, for the fMRI study we probed the modulation of accuracy by confidence, the proportion of high confidence false percepts on grating-absent trials, and the proportion of cue congruent responses. Participants’ orientation responses were also explored with a logistical regression model, but without stimulus contrast as a predictor as this was not varied for the purposes of the fMRI experiment.

## Notes

### Competing Interest Statement

The authors have declared no competing interest.

### Summary of Updates

We added control analyses to account for a possible contribution of the noise patches to the bottom-up false percepts. We also corrected a number of small typos.

## References

1. Weil, R. S. et al. Visual dysfunction in Parkinson’s disease. Brain 139, 2827–2843 (2016).

2. Powers, A. R., Kelley, M. & Corlett, P. R. Hallucinations as Top-Down Effects on Perception. Biol. Psychiatry Cogn. Neurosci. Neuroimaging 1, 393–400 (2016).

3. Corlett, P. R. et al. Hallucinations and Strong Priors. Trends Cogn. Sci. 23, 114–127 (2019).

4. Reichert, D. P., Seriès, P. & Storkey, A. J. Charles Bonnet Syndrome: Evidence for a Generative Model in the Cortex? PLoS Comput. Biol. 9, e1003134 (2013).

5. Sterzer, P. et al. The Predictive Coding Account of Psychosis. Biol. Psychiatry 84, 634–643 (2018).

6. Friston, K. The free-energy principle: a rough guide to the brain? Trends Cogn. Sci. 13, 293–301 (2009).

7. Bastos, A. M. et al. Canonical Microcircuits for Predictive Coding. Neuron 76, 695–711 (2012).

8. Haarsma, J., Kok, P. & Browning, M. The promise of layer-specific neuroimaging for testing predictive coding theories of psychosis. Schizophr. Res. 245, 68–76 (2022).

9. de Lange, F. P., Heilbron, M. & Kok, P. How Do Expectations Shape Perception? Trends Cogn. Sci. 22, 764–779 (2018).

10. Powers, A. R., Mathys, C. & Corlett, P. R. Pavlovian conditioning–induced hallucinations result from overweighting of perceptual priors. Science 357, 596–600 (2017).

11. Kafadar, E. et al. Modeling perception and behavior in individuals at clinical high risk for psychosis: Support for the predictive processing framework. Schizophr. Res. 226, 167– 175 (2020).

12. Cassidy, C. M. et al. A Perceptual Inference Mechanism for Hallucinations Linked to Striatal Dopamine. Curr. Biol. 28, 503–514.e4 (2018).

13. Haarsma, J. et al. Influence of prior beliefs on perception in early psychosis: Effects of illness stage and hierarchical level of belief. J. Abnorm. Psychol. 129, 581–598 (2020).

14. Teufel, C. et al. Shift toward prior knowledge confers a perceptual advantage in early psychosis and psychosis-prone healthy individuals. Proc. Natl. Acad. Sci. 112, 13401– 13406 (2015).

15. Schmack, K. et al. Delusions and the Role of Beliefs in Perceptual Inference. J. Neurosci. 33, 13701–13712 (2013).

16. Schmack, K., Schnack, A., Priller, J. & Sterzer, P. Perceptual instability in schizophrenia: Probing predictive coding accounts of delusions with ambiguous stimuli. Schizophr. Res. Cogn. 2, 72–77 (2015).

17. Schmack, K., Bosc, M., Ott, T., Sturgill, J. F. & Kepecs, A. Striatal dopamine mediates hallucination-like perception in mice. Science 372, eabf4740 (2021).

18. Stuke, H., Kress, E., Weilnhammer, V. A., Sterzer, P. & Schmack, K. Overly Strong Priors for Socially Meaningful Visual Signals Are Linked to Psychosis Proneness in Healthy Individuals. Front. Psychol. 12, 583637 (2021).

19. Hahamy, A., Wilf, M., Rosin, B., Behrmann, M. & Malach, R. How do the blind ‘see’? The role of spontaneous brain activity in self-generated perception. Brain 144, 340–353 (2021).

20. Burke, W. The neural basis of Charles Bonnet hallucinations: a hypothesis. J. Neurol. Neurosurg. Psychiatry 73, 535–541 (2002).

21. Desai, N. S., Rutherford, L. C. & Turrigiano, G. G. Plasticity in the intrinsic excitability of cortical pyramidal neurons. Nat. Neurosci. 2, 515–520 (1999).

22. Painter, D. R., Dwyer, M. F., Kamke, M. R. & Mattingley, J. B. Stimulus-Driven Cortical Hyperexcitability in Individuals with Charles Bonnet Hallucinations. Curr. Biol. 28, 3475–3480.e3 (2018).

23. Denève, S. & Jardri, R. Circular inference: mistaken belief, misplaced trust. Curr. Opin. Behav. Sci. 11, 40–48 (2016).

24. Jardri, R., Duverne, S., Litvinova, A. S. & Denève, S. Experimental evidence for circular inference in schizophrenia. Nat. Commun. 8, 14218 (2017).

25. Teufel, C., Kingdon, A., Ingram, J. N., Wolpert, D. M. & Fletcher, P. C. Deficits in sensory prediction are related to delusional ideation in healthy individuals. Neuropsychologia 48, 4169–4172 (2010).

26. Notredame, C.-E., Pins, D., Deneve, S. & Jardri, R. What visual illusions teach us about schizophrenia. Front. Integr. Neurosci. 8, (2014).

27. Weilnhammer, V. et al. Psychotic Experiences in Schizophrenia and Sensitivity to Sensory Evidence. Schizophr. Bull. 46, 927–936 (2020).

28. Boly, M. et al. Baseline brain activity fluctuations predict somatosensory perception in humans. Proc. Natl. Acad. Sci. 104, 12187–12192 (2007).

29. Busch, N. A., Dubois, J. & VanRullen, R. The Phase of Ongoing EEG Oscillations Predicts Visual Perception. J. Neurosci. 29, 7869–7876 (2009).

30. Wyart, V. & Tallon-Baudry, C. How Ongoing Fluctuations in Human Visual Cortex Predict Perceptual Awareness: Baseline Shift versus Decision Bias. J. Neurosci. 29, 8715– 8725 (2009).

31. Pajani, A., Kok, P., Kouider, S. & de Lange, F. P. Spontaneous Activity Patterns in Primary Visual Cortex Predispose to Visual Hallucinations. J. Neurosci. 35, 12947–12953 (2015).

32. Podvalny, E., Flounders, M. W., King, L. E., Holroyd, T. & He, B. J. A dual role of prestimulus spontaneous neural activity in visual object recognition. Nat. Commun. 10, 3910 (2019).

33. Felleman, D. J. & Van Essen, D. C. Distributed Hierarchical Processing in the Primate Cerebral Cortex. Cereb. Cortex 1, 1–47 (1991).

34. Harris, K. D. & Mrsic-Flogel, T. D. Cortical connectivity and sensory coding. Nature 503, 51–58 (2013).

35. Muckli, L. et al. Contextual Feedback to Superficial Layers of V1. Curr. Biol. 25, 2690– 2695 (2015).

36. Kok, P., Bains, L. J., van Mourik, T., Norris, D. G. & de Lange, F. P. Selective Activation of the Deep Layers of the Human Primary Visual Cortex by Top-Down Feedback. Curr. Biol. 26, 371–376 (2016).

37. Lawrence, S. J. D., Formisano, E., Muckli, L. & de Lange, F. P. Laminar fMRI: Applications for cognitive neuroscience. NeuroImage 197, 785–791 (2019).

38. Stephan, K. E. et al. Laminar fMRI and computational theories of brain function. NeuroImage 197, 699–706 (2019).

39. Self, M. W., van Kerkoerle, T., Goebel, R. & Roelfsema, P. R. Benchmarking laminar fMRI: Neuronal spiking and synaptic activity during top-down and bottom-up processing in the different layers of cortex. NeuroImage 197, 806–817 (2019).

40. Jia, K. et al. Recurrent Processing Drives Perceptual Plasticity. Curr. Biol. 30, 4177–4187.e4 (2020).

41. Ng, A. K. T. et al. Ultra-High-Field Neuroimaging Reveals Fine-Scale Processing for 3D Perception. J. Neurosci. 41, 8362–8374 (2021).

42. Aitken, F. et al. Prior expectations evoke stimulus-specific activity in the deep layers of the primary visual cortex. PLOS Biol. 18, e3001023 (2020).

43. Sheldon, A. D. et al. Perceptual pathways to hallucinogenesis. Schizophr. Res. (2022).

44. Lawrence, S. J., Norris, D. G. & de Lange, F. P. Dissociable laminar profiles of concurrent bottom-up and top-down modulation in the human visual cortex. eLife 8, e44422 (2019).

45. Lawrence, S. J. D. et al. Laminar Organization of Working Memory Signals in Human Visual Cortex. Curr. Biol. 28, 3435–3440.e4 (2018).

46. Bell, V., Halligan, P. W. & Ellis, H. D. The Cardiff Anomalous Perceptions Scale (CAPS): A New Validated Measure of Anomalous Perceptual Experience. Schizophr. Bull. 32, 366– 377 (2006).

47. Ress, D. & Heeger, D. J. Neuronal correlates of perception in early visual cortex. Nat. Neurosci. 6, 414–420 (2003).

48. Leptourgos, P., Bouttier, V., Denève, S. & Jardri, R. From hallucinations to synaesthesia: A circular inference account of unimodal and multimodal erroneous percepts in clinical and drug-induced psychosis. Neurosci. Biobehav. Rev. 135, 104593 (2022).

49. van Kerkoerle, T., Self, M. W. & Roelfsema, P. R. Layer-specificity in the effects of attention and working memory on activity in primary visual cortex. Nat. Commun. 8, 13804 (2017).

50. Dijkstra, N., Kok, P. & Fleming, S. M. Perceptual reality monitoring: Neural mechanisms dissociating imagination from reality. Neurosci. Biobehav. Rev. 135, 104557 (2022).

51. Adams, R. A., Stephan, K. E., Brown, H. R., Frith, C. D. & Friston, K. J. The Computational Anatomy of Psychosis. Front. Psychiatry 4, (2013).

52. Landgraf, S. & Osterheider, M. “To see or not to see: that is the question.” The “Protection-Against-Schizophrenia” (PaSZ) model: evidence from congenital blindness and visuo-cognitive aberrations. Front. Psychol. 4, (2013).

53. ffytche, D. H. et al. The anatomy of conscious vision: an fMRI study of visual hallucinations. Nat. Neurosci. 1, 738–742 (1998).

54. Zarkali, A. et al. Increased weighting on prior knowledge in Lewy body-associated visual hallucinations. Brain Commun. 1, fcz007 (2019).

55. Pagonabarraga, J. et al. Minor hallucinations occur in drug-naive Parkinson’s disease patients, even from the premotor phase: Minor Hallucinations in Untreated PD Patients. Mov. Disord. 31, 45–52 (2016).

56. Mocellin, R., Walterfang, M. & Velakoulis, D. Neuropsychiatry of Complex Visual Hallucinations. 10 (2006).

57. D’Antonio, F. et al. Visual hallucinations in Lewy body disease: pathophysiological insights from phenomenology. J. Neurol. 269, 3636–3652 (2022).

58. Mazor, M., Friston, K. J. & Fleming, S. M. Distinct neural contributions to metacognition for detecting, but not discriminating visual stimuli. eLife 9, e53900 (2020).

59. Samaha, J., Iemi, L., Haegens, S. & Busch, N. A. Spontaneous Brain Oscillations and Perceptual Decision-Making. Trends Cogn. Sci. 24, 639–653 (2020).

60. Spruston, N. Pyramidal neurons: dendritic structure and synaptic integration. Nat. Rev. Neurosci. 9, 206–221 (2008).

61. Larkum, M. A cellular mechanism for cortical associations: an organizing principle for the cerebral cortex. Trends Neurosci. 36, 141–151 (2013).

62. Binzegger, T. A Quantitative Map of the Circuit of Cat Primary Visual Cortex. J. Neurosci. 24, 8441–8453 (2004).

63. Kim, J., Matney, C. J., Blankenship, A., Hestrin, S. & Brown, S. P. Layer 6 Corticothalamic Neurons Activate a Cortical Output Layer, Layer 5a. J. Neurosci. 34, 9656– 9664 (2014).

64. Antic, S. D., Zhou, W.-L., Moore, A. R., Short, S. M. & Ikonomu, K. D. The decade of the dendritic NMDA spike. J. Neurosci. Res. 88, 2991–3001 (2010).

65. Major, G., Larkum, M. E. & Schiller, J. Active Properties of Neocortical Pyramidal Neuron Dendrites. Annu. Rev. Neurosci. 36, 1–24 (2013).

66. Hawkins, J. & Ahmad, S. Why Neurons Have Thousands of Synapses, a Theory of Sequence Memory in Neocortex. Front. Neural Circuits 10, (2016).

67. Hawkins, J., Ahmad, S. & Cui, Y. A Theory of How Columns in the Neocortex Enable Learning the Structure of the World. Front. Neural Circuits 11, 81 (2017).

68. Jardri, R. et al. Are Hallucinations Due to an Imbalance Between Excitatory and Inhibitory Influences on the Brain? Schizophr. Bull. 42, 1124–1134 (2016).

69. Meijs, E. L., Slagter, H. A., de Lange, F. P. & van Gaal, S. Dynamic Interactions between Top–Down Expectations and Conscious Awareness. J. Neurosci. 38, 2318–2327 (2018).

70. Alilović, J., Slagter, H. A. & van Gaal, S. Subjective visibility report is facilitated by conscious predictions only. Conscious. Cogn. 87, 103048 (2021).

71. Kok, P., Brouwer, G. J., van Gerven, M. A. J. & de Lange, F. P. Prior Expectations Bias Sensory Representations in Visual Cortex. J. Neurosci. 33, 16275–16284 (2013).

72. Chalk, M., Seitz, A. R. & Series, P. Rapidly learned stimulus expectations alter perception of motion. J. Vis. 10, 2–2 (2010).

73. Aitken, F. & Kok, P. Hippocampal representations switch from errors to predictions during acquisition of predictive associations. 31.

74. Aru, J., Suzuki, M., Rutiku, R., Larkum, M. E. & Bachmann, T. Coupling the State and Contents of Consciousness. Front. Syst. Neurosci. 13, 43 (2019).

75. Takahashi, N. et al. Active dendritic currents gate descending cortical outputs in perception. Nat. Neurosci. 23, 1277–1285 (2020).

76. Kok, P., Failing, M. F. & de Lange, F. P. Prior Expectations Evoke Stimulus Templates in the Primary Visual Cortex. J. Cogn. Neurosci. 26, 1546–1554 (2014).

77. Foster, K. H., Gaska, J. P., Nagler, M. & Pollen, D. A. Spatial and temporal frequency selectivity of neurones in visual cortical areas V1 and V2 of the macaque monkey. J. Physiol. 365, 331–363 (1985).

78. Peters, E., Joseph, S., Day, S. & Garety, P. Measuring Delusional Ideation: The 21-Item Peters et al. Delusions Inventory (PDI). Schizophr. Bull. 30, 1005–1022 (2004).

79. Wyart, V., Nobre, A. C. & Summerfield, C. Dissociable prior influences of signal probability and relevance on visual contrast sensitivity. Proc. Natl. Acad. Sci. 109, 3593– 3598 (2012).

80. Anwyl-Irvine, A. L., Massonnié, J., Flitton, A., Kirkham, N. & Evershed, J. K. Gorilla in our midst: An online behavioral experiment builder. Behav. Res. Methods 52, 388–407 (2020).

81. van Mourik, T., van der Eerden, J. P. J. M., Bazin, P.-L. & Norris, D. G. Laminar signal extraction over extended cortical areas by means of a spatial GLM. PLOS ONE 14, e0212493 (2019).

82. Kleinnijenhuis, M. et al. Diffusion tensor characteristics of gyrencephaly using high resolution diffusion MRI in vivo at 7T. NeuroImage 109, 378–387 (2015).

83. Waehnert, M. D. et al. Anatomically motivated modeling of cortical laminae. NeuroImage 93, 210–220 (2014).

84. de Sousa, A. A. et al. Comparative Cytoarchitectural Analyses of Striate and Extrastriate Areas in Hominoids. Cereb. Cortex 20, 966–981 (2010).

85. Cousineau, D. Confidence intervals in within-subject designs: A simpler solution to Loftus and Masson’s method. Tutor. Quant. Methods Psychol. 1, 42–45 (2005).

86. Morey, R. D. Confidence Intervals from Normalized Data: A correction to Cousineau (2005). Tutor. Quant. Methods Psychol. 4, 61–64 (2008).

