## Supplementary Material for "Perceptual expectations and false percepts generate stimulus-specific activity in distinct layers of the early visual cortex"

*Layer-specific effects of stimulus presentation and functional localiser*

As reported in the main text, a cross-validated analysis of orientation-specific BOLD signals within the functional localiser revealed stronger orientation-specific effects in V2 than V1 across all layers (main effect of ROI: F{1,23}=23.56, *p*<.001, all layers *p*<.01)*;* Supplementary Fig. 2a). Further, the noisy gratings on grating-present trials evoked significant orientation-specific activity in the superficial layers of V2 (T{24}=2.00, *p*=.028) but not the other layers (both *p*>.1; Supplementary Fig. 2b). Confidence did not significantly affect orientation-specific activity on grating-present trials (F{1,24}=3.18, *p*=.081).

For visualisation purposes, we plot the orientation-specific activity on grating-absent trials across all four different confidence levels, rather than dividing them into high and low confidence, for false percept effects (Supplementary Fig. 2d). Since there was considerable variability in the number of different confidence responses across the different conditions, not every condition will contain the same number of subjects, and we therefore did not perform any statistical tests on these effects. On grating-absent trials (Supplementary Fig. 2d), this revealed, at least numerically, stronger activity in the middle layers for confidence levels in the higher range (3 and 4), compared to those in the lower range (1 and 2), in line with the core results demonstrating the middle layers to be involved in hallucinations (Supplementary Fig. 2d).

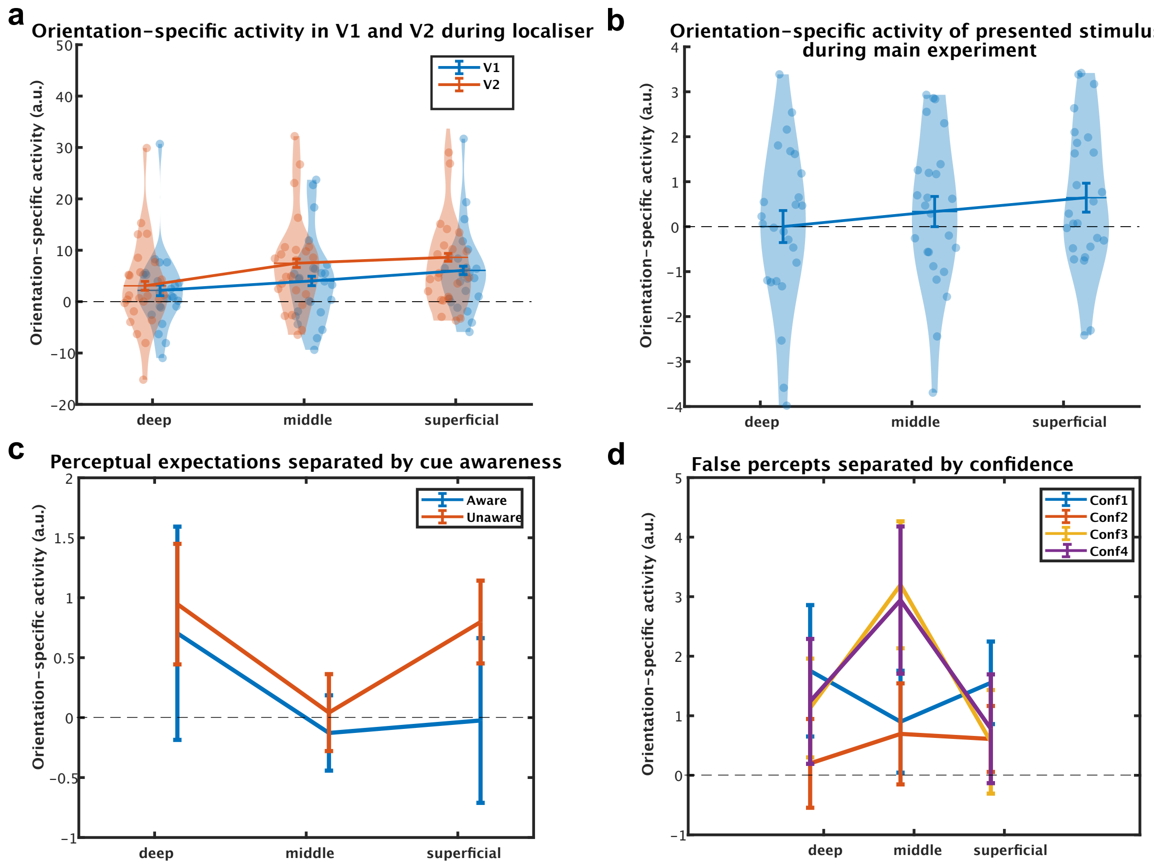


***Supplementary Fig. 1 | Supplementary visualisations of layer- specific effects. a**, Stimulus-specific activity in V2 was stronger than in V1. **b**, Stimulus-specific activity in the superficial layers evoked by presented gratings during the main*

*experiment (grating-present trials).* ***c****, Perceptual expectation- induced activity, separated by cue awareness.* ***d****, False percept-induced activity for all four confidence levels, showing an increase in middle layer activity with increasing confidence. Bars represent within-subject error bars, except in figure d where they represent standard error of the mean.*

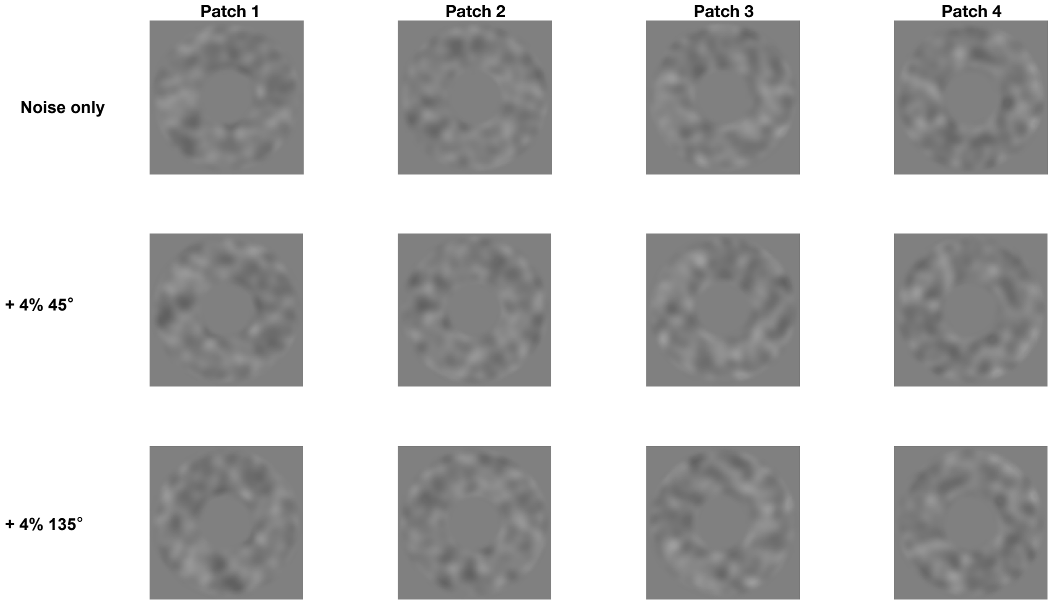


***Supplementary Fig. 2 | Visual stimuli used in the present experiment.*** *Top row: Four 20% contrast noise patches with flat orientation energy spectra were generated to avoid stimulus-driven orientation signals on grating-absent trials (also see Supplementary Fig. 5). Middle row: 45° grating-present trials were created by adding 4% contrast 45° gratings to the four noise patches. Bottom row: Likewise, 135° grating-present trials were created by adding 4% contrast 135° gratings to the four noise patches.*

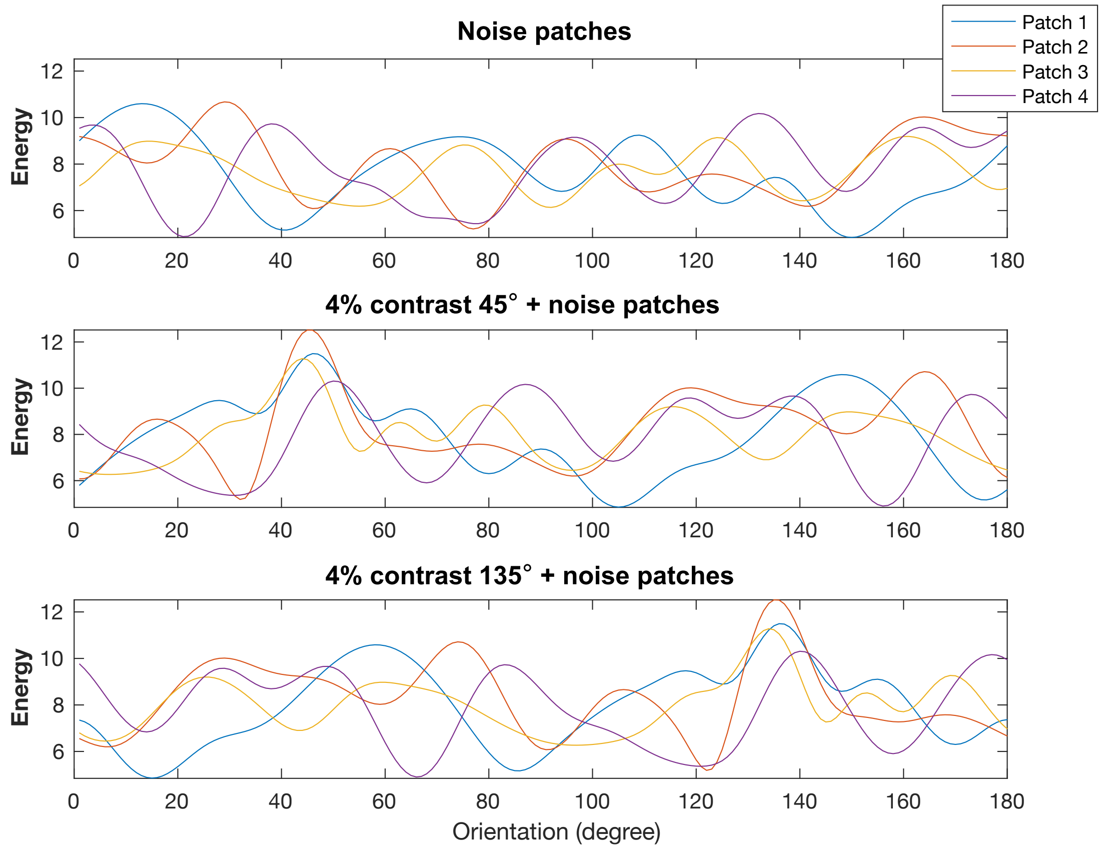


***Supplementary Fig. 3 | Orientation filter output.*** *The output of the orientation filters when applied to the noise patches (top row), as well as the 45° grating-present trials (middle row), and the 135° grating-present trials (bottom row). Notice that the orientation energy for grating-present trials peaks around 45° and 135°, respectively, as expected.*

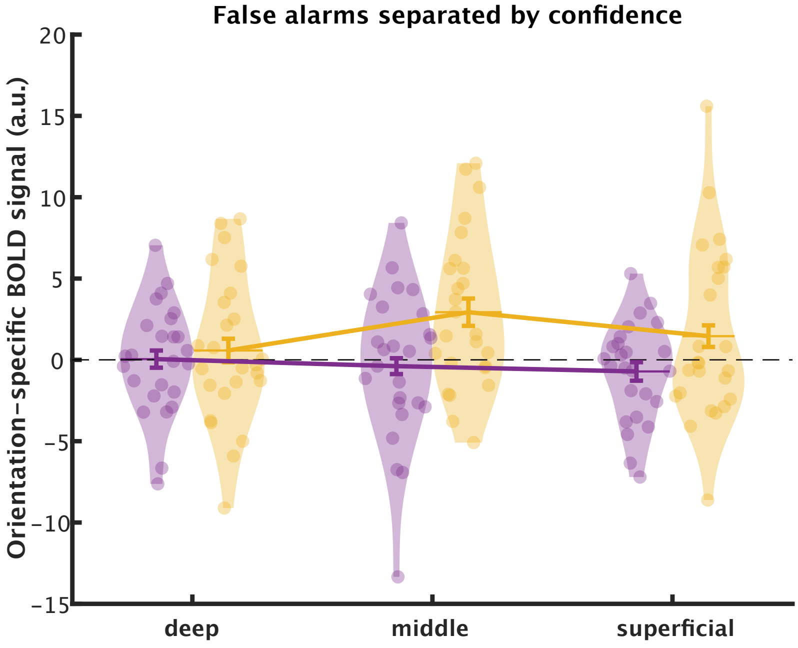


***Supplementary Fig. 4 | Analysis controlling for noise patches, showing orientation-specific BOLD activity in the cortical layers of V2.*** *In order to control for the possibility that the noise patches themselves contributed to the orientation-specific BOLD responses in the middle layers for high confidence false percepts, we modelled*

*the effects of high and low confidence 45° and 135° false percepts separately for each of the four noise patches, to ensure they contributed equally to the orientation-specific effects. This analysis replicated our main finding that high confidence false percepts are reflected in the middle layers.*

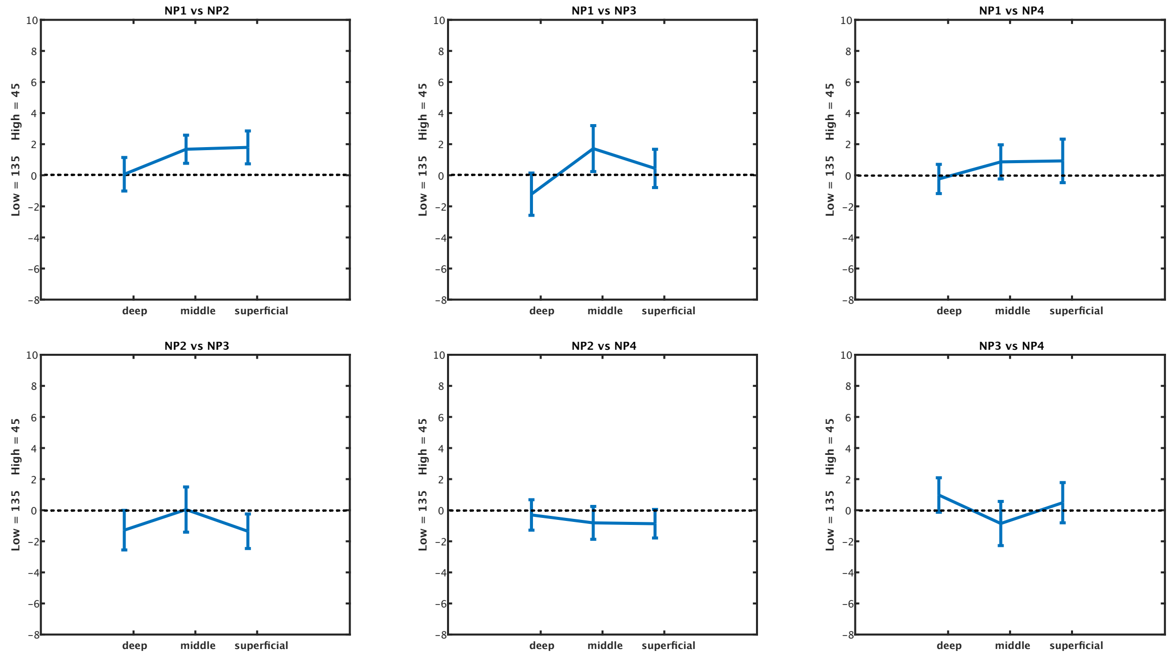


***Supplementary Fig. 5 | Differences in stimulus-specific BOLD activity between noise patches.*** *Here we estimated for each noise patch the degree to which each induces stimulus-specific activity by subtracting the activity profile in 45° and 135°-preferring voxels. We then tested for all six possible pairwise comparisons of noise patches whether there was a difference in orientation-specific activity. Note that noise patches two and three were more often identified as 45°, and noise patch four more often as 135°. Therefore, if our main results were driven by the noise patches themselves, we would expect to see the largest differences in orientation-specific activity when comparing noise patches two and four (bottom-middle) and three and four (bottom-right). However, no significant differences were found for any of the pairwise comparison (all p>.05, uncorrected), rendering an explanation of our effects being driven by bottom-up signals present in the noise patches unlikely. NP = Noise patch.*

### *Increase in stimulus contrast resulted in increased confidence and accuracy in the online experiment*

We explored whether the high confidence false alarm responses that are related to middle layer activity in V2 predict everyday hallucination severity. We conducted a large online study in which questionnaires measuring the prevalence of hallucinatory percepts in daily life were collected alongside the false percept task. The false percept task used here was virtually identical to the one used in the scanner (with slight variations in practice procedure and trial counts). One important difference was the introduction of three (rather than one) contrast levels on the grating-present trials, to enable estimates of sensory precision, i.e., how task accuracy depended on evidence quality. Specifically, a base-level contrast value was selected for each participant based on their performance during the instruction phase. This base-contrast was used during the main experiment along with gratings with 1% higher or lower contrast as well as absent trials. In the online study 22 out of the 100 participants became aware of the cue. Confidence increased with contrast (0%, base-1%, base, base+1%) ($F_{\{2,297\}}$=133.8, *p*<.001; all post-hoc tests *p*<.001) (Supplementary Fig. 6a), but there was no effect of cue validity on confidence ($F_{\{1,99\}}$=1.952, *p*=.17), nor was there an interaction between cue validity and stimulus contrast (F{1,99}=0.87, *p*=.42). Accuracy increased with contrast (F{2,196}=49.3, *p*<.001), and was lower when the expectation cue was invalid (F{1,98}=20.50, *p*<.001). The effect of the expectation cues interacted with participants' awareness of the meaning of the cues (F{1,98}=10.7, p=.001), such that the cue effects were stronger in those who were aware of the cue's meaning (Supplementary Fig. 6b). Furthermore, the expectation cues influenced participants choice behaviour on grating-absent trials (T{99}=2.97, *p*=.004). This was driven by those who became aware of the cue meaning (N=22, 58.3% false percepts congruent with the cue), who were significantly more influenced by the cues than those who were not aware of their meaning (N=78, 51.6% false percepts congruent with the cue; T{98}=2.83, *p*=.006). Those unaware of the cues' meaning only showed trend-level responses in line with cue (T{77}=1.87,*p*=.065), while those aware showed a significant effect of cue (T{21}=2.41, *p*=.025). These findings are similar to those of the fMRI study, where the effect of cue was also driven by those aware of the cues' purpose. Further, accuracy and confidence increased with grating contrast as expected. We modelled choice behaviour on grating-absent and grating-present trials using a logistic regression model. This revealed that responses on grating-present trials were driven by the interaction of the current stimulus and contrast (hereafter referred to as sensory precision) (T{99}=14.11, *p*<.001), previous response (T{99}=8.93, *p*<.001) and current stimulus (T{99}=8.07, *p*<.001) (Supplementary Fig. 6c). Conversely, responses on grating-absent trials were driven by previous responses (T{99}=2.34, *p*=.021) (Supplementary Fig. 6d).

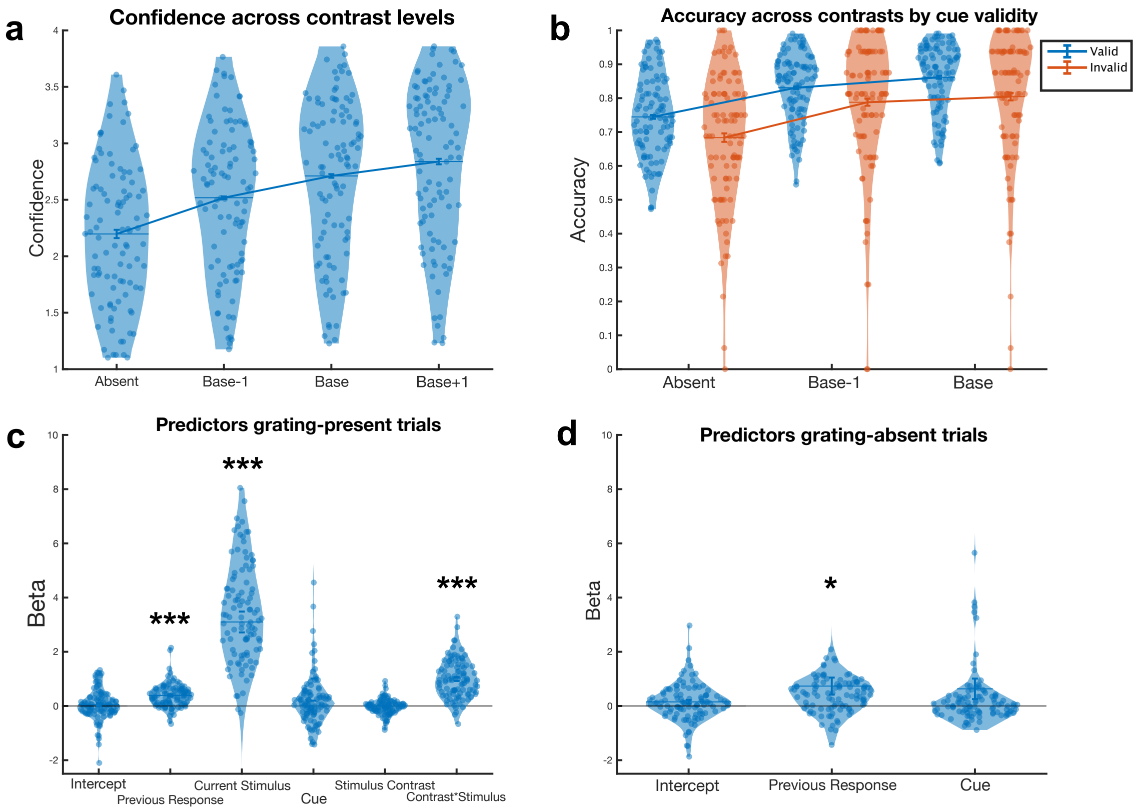


***Supplementary Fig. 6 | Behavioural results from online study.*** ***a**, Confidence increased across contrast levels.* ***b**, Accuracy increased across contrast levels and was higher for valid trials, but this was driven by those aware of cues.* ***c**, Previous response, current stimulus, and the interaction between current stimulus and contrast were significant predictors for grating-present trials.* ***d**, Previous responses predicted grating-absent trial responses. Bars represent standard error of the mean. * = p<.05 *** = p<.001.*
